# polyDFEv2.0: Testing for invariance of the distribution of fitness effects within and across species

**DOI:** 10.1101/363887

**Authors:** Paula Tataru, Thomas Bataillon

## Abstract

Distributions of fitness effects (DFE) of mutations can be inferred from site frequency spectrum (SFS) data. There is mounting interest to determine whether distinct genomic regions and/or species share a common DFE, or whether evidence exists for differences among them. polyDFEv2.0 fits multiple SFS datasets at once and provides likelihood ratio tests for DFE invariance across datasets. Simulations show that testing for DFE invariance across genomic regions within a species requires models accounting for heterogeneous genealogical histories underlying SFS data in these regions. Not accounting for these heterogeneities will result in the spurious detection of DFE differences.

## Introduction

Levels of purifying and positive selection vary throughout the genome and a powerful way to study such variation is by inferring the distribution of fitness effects (DFE) for different genomic regions that are a priori expected to undergo distinct selective pressures (Gronau et al. 2013; Racimo and Schraiber 2014). Similarly, there is renewed interest in understanding what genomic, demographic or ecological factors explain differences in genome-wide polymorphism patterns across species (Ellegren and Galtier 2016; Chen et al. 2017; Huber et al. 2017). The site frequency spectrum (SFS) contains information to infer the DFE of new mutations. Existing methods infer the DFE while accounting for demography and other sources of distortion in the SFS (Schneider et al. 2011; Kousathanas and Keightley 2013; Galtier 2016; Kim et al. 2017; Tataru et al. 2017; Barton et al. 2018). Yet current methods assume that a mutation’s fitness is drawn from a single common DFE and are therefore not well suited to determine whether distinct gene categories, genomic regions and/or species share invariant DFEs, or whether there are genuine differences between such categories.

Here we present a new method, polyDFEv2.0, testing for DFEs invariance across datasets, be it distinct genomic regions within species, or different species. Simulations demonstrate that the method guards against excessive type I error while retaining substantial power to detect differences across datasets.

## Results and Discussion

polyDFE implements a likelihood framework that allows fitting simultaneously DFE parameters, nuisance parameters that account for distortions in the SFS data induced by linkage and demography and errors when polarizing the SFS data (Tataru et al. 2017). Here, we extend polyDFE for fitting multiple datasets. Any fitted parameter can be constrained to be shared (invariant) across datasets or fitted independently for each dataset (models M_1_ to M_4_, Table 1). Likelihood ratio tests (LRT) are used to determine if the datasets have different DFEs.

**Table 1:** Models for detecting heterogeneity in DFEs across datasets fitted in polyDFE

| Models for jointly fitting distinct SFS datasets | Model parameters fitted |  |  |  |
| --- | --- | --- | --- | --- |
|  | DFE parameters | Demography | Mutation rate | Polarization error |
| $M_1$ | shared | shared | shared | shared |
| $M_2$ | independent | shared | shared | shared |
| $M_3$ | shared | independent | independent | independent |
| $M_4$ | independent | independent | independent | independent |

When testing for invariance in DFEs across species, we use LRTs comparing the fit of M_3_ and M_4_, as we expect demography and scaled mutation rates to vary between species (Table 1). Simulations (Supplementary Methods) show that comparing M_3_ and M_4_ provides a reliable test for the null hypothesis of DFE invariance: no inflation in type 1 error under H_0_ and good power to detect differences in strength of purifying and positive selection (Figure S1).

When analyzing SFS datasets from distinct regions or gene ontologies within one species, we investigated the type I error and power of two LRTs: M_1_ versus M_2_ and M_3_ versus M_4_. Comparing models M_1_ and M_2_ amounts to assuming that all genomic regions share the same nuisance parameters. Simulations show that M_1_ might not be a proper null model for DFE invariance within species (Figures S1 and S2). This is because even under DFE invariance, differences in coalescent histories between genomic regions and SNP polarization error rates have to be accounted for. Even when fitting different SFS datasets within a single species, we recommend fitting models with nuisance parameters for each SFS. Using models that allow for both differences in DFE and nuisance parameters (M_3_ and M_4_) allows to control for type I error while retaining substantial power to detect differences in DFE (Figure S1).

We re-analyzed a chimpanzee dataset (Bataillon et al. 2015) and tested for DFEs invariance across two subspecies (central vs. eastern chimpanzee) and between autosomal vs. X-linked regions. We found autosomal DFEs to be different between subspecies. Central chimpanzees exhibit a higher effective size resulting in overall more strongly deleterious mutations (Figure S3A). No significant differences are found when the comparison involved X-linked regions (Figure S3B and C). While genuine differences in DFE probably exist, simulations show that the amount of data available on the X chromosome (0.89Mb of X linked sites vs 20Mb of autosomes) entails substantial loss of power. The amount of data available within each SFS dataset can limit drastically the statistical power to detect these differences (Figure S4).

## Conclusion

polyDFEv2.0 provides reliable tests for DFE invariance. It is flexible enough to also test more specific hypothesis regarding the nature of the differences across DFEs: e.g. did the shape of the DFE change and/or the proportion of beneficial mutations change across DFEs? Using models that are flexible enough to account jointly for differences in nuisance parameters across categories/species is of paramount importance to make reliable tests for DFE invariance across and within species.

## Supplementary Material

polyDFEv2.0 is available as source code and compiled binaries under a GNU General Public License v3.0 from https://github.com/paula-tataru/polyDFE.

Supplementary Figures S1 - S4, Supplementary table S1 and Supplementary Material are available online.

## Acknowledgments

This work has been supported by the European Research Council under the European Union’s Seventh Framework Program (FP7/20072013, ERC grant number 311341). We thank D. Castellano, M. Hartfield, and E. Lucotte for comments on previous drafts of the manuscript.

## Supplementary information

### SFS data simulation setups

To investigate the type I error rate and power in detecting if two datasets share the same DFE or not, we relied on simulated SFS data obtained from exome-like regions using SFS_CODE (Hernandez 2008). Among the 18 DFE scenarios used to validate the statistical performance of the polyDFE method (Tataru et al. 2017), we explored here data generated under 4 possible types of DFEs (described in Table S1). Some DFEs comprise only deleterious mutation and vary intensity of purifying selection as measured by ***S***_d_, the mean scaled effects of mutations (DelHSD, DelMSD, Table S1). Two other DFEs contain a fraction ***p***_b_ of beneficial mutations with different mean scaled positive selective coefficient ***S***_b_ resulting in different expected amounts of adaptive evolution as measured by the parameter α (MAMSD, HAHSB, Table S1). For each DFE type, we simulated 100 replicate datasets comprising SFS data containing 10.8 Mb of exonic sites in 10 diploid individuals (20 haplotypes). Each dataset was simulated using a scaled mutation rate of 0.001. The simulated data considered here do not contain misidentification of the ancestral state or demography (i.e. population size was assumed to be constant).

**Table S1.** Propeties of DFEs assumed when simulating SFS datsets

| | DFE type | DFE abbreviation | $S_d$ | $b$ | $p_b$ | $S_b$ | $\alpha$ |
| --- | --- | --- | --- | --- | --- | --- | --- |
| deleterious DFEs | high $S_d$ | DelHSD | -10 000 | 0.4 | 0 | - | 0 |
| | medium $S_d$ | DelMSD | -400 | 0.4 | 0 | - | 0 |
| full DFEs | medium $\alpha$, medium $S_d$ | MAMSD | -400 | 0.4 | 0.02 | 4 | 0.53 |
| | high $\alpha$, high $S_b$ | HAHSB | -400 | 0.4 | 0.02 | 16 | 0.81 |

Note The DFEs consist of a mixture between a gamma and exponential distributions: a new mutation is deleterious with probability $1 - p_b$ , and has a scaled selection coefficient drawn from a reflected gamma distribution with mean $S_d < 0$ and shape $b$ ; a new mutation is beneficial with probability $p_b$ , and has a (scaled) selection coefficient drawn from an exponential distribution with mean $S_b > 0$ .

### Details of inference on simulated data

The inference was performed for each simulated dataset under models M_1_-M_4_ presented in Table 1 in the main text. polyDFEv2.0 was ran on two simulated datasets, where these datasets were either simulated using the same underlying DFE (when investigating type I error) or using two different DFEs (for investigating the power to detect differences in DFEs, see below). When fitting the data using polyDFEv2.0, the ancestral misidentification error ***ε*** was fixed to 0 and not estimated, while the demography nuisance parameters (*r_i_*s) were estimated, as it has been shown that fitting these accounts for SFS distortions induced by demographic history (relative to the SFS expectation in a stable population) but also the presence of linkage in the data (Tataru et al. 2017). The scaled mutation rate parameter was assumed constant across the data. To ensure that the likelihood function was reliably maximized, we performed 10 runs of the BFGS algorithm, with randomly starting values for each dataset analysed. We also provided the simulated values of the parameters to polyDFE as an additional starting point, to ensure that a failure in finding the simulated values as the true optimum was not caused by a failure of finding the global optimum.

### Evaluating the type 1 error and power of likelihood ratio tests for DFE invariance

polyDFE’s statistical performance for estimating DFE parameters has already been validated using datasets simulated under a broad range of DFE types and demographic scenarios (Tataru et al. 2017), including the four DFE types listed in Table S1. Therefore, we focus on the performance of two likelihood ratio tests (LRT): LRT comparing models M_1_ vs M_2_ and M_3_ vs M_4_. The percentage of p-values that are below the 5% threshold then gives the type I error (when the 2 input datasets were simulated using the same underlying DFE) and the power (when the 2 input datasets were simulated using different underlying DFEs). When performing LRT, the distribution of the LRT statistic obtained from fitting data under the null model (invariant DFE) is asymptotically approximated by a χ^2^ distribution (this approximation is expected to be valid when the sample size is large). When using datasets of approximately 10Mb of sites each, we assumed a χ^2^ distribution with 4 degrees of freedom to calculate the p-value for the LRT (Figure S1) for both model comparisons (models M_1_ vs M_2_ and M_3_ vs M_4_).

However, when smaller datasets are analysed, the empirical null distribution of the LRT statistic is no longer reliably approximated by a χ^2^ distribution (Figure S2). Therefore, to investigate how power changes as a function of the size of the dataset (Figure S4), we relied on the empirical (simulated) null distribution of the LRT statistic to obtain the p-values. To avoid using the χ^2^ distribution when testing for a DFE invariance among datasets of small size, one should ideally simulate multiple datasets under the inferred null model, infer the parameters for these datasets using the null and alternative models and obtain an empirical null distribution for the LRT statistic. The simulation can be performed directly from polyDFE.

When comparing models M_1_ and M_2_, the empirical null distribution of the LRT statistic is badly approximated by the χ^2^ distribution (Figure S2), even when the amount data is large (10Mb), which leads to a high type I error (Figure S1). However, the χ^2^approximation is much better when comparing models M_3_ and M_4_. This suggests that model M_1_ may not be a proper null model for determining if the DFE is shared or not between datasets.

### Details of inference on the chimpanzee datasets

We analysed a chimpanzee dataset and test: (1) if the central and eastern chimpanzee subspecies shared the same DFE, (2) if autosomes and X chromosome share the same DFE in each chimpanzee subspecies. To do so, we fitted the data in polyDFEv2.0 using models M_3_ and M_4_ (Figure S3). The inference was performed on the SFS data only (divergence counts to an outgroup were not fitted) and SFS data was fitted using a DFE model comprising both deleterious and beneficial mutations, and both an ancestral SNP misidentification error *ε* and distortion parameters (*r_i_*s) were estimated. Mutation was assumed constant across the data. To ensure that the likelihood function was reliably maximized, we performed 10 runs of the BFGS algorithm, with randomly starting values.

---

**Figure S1.**
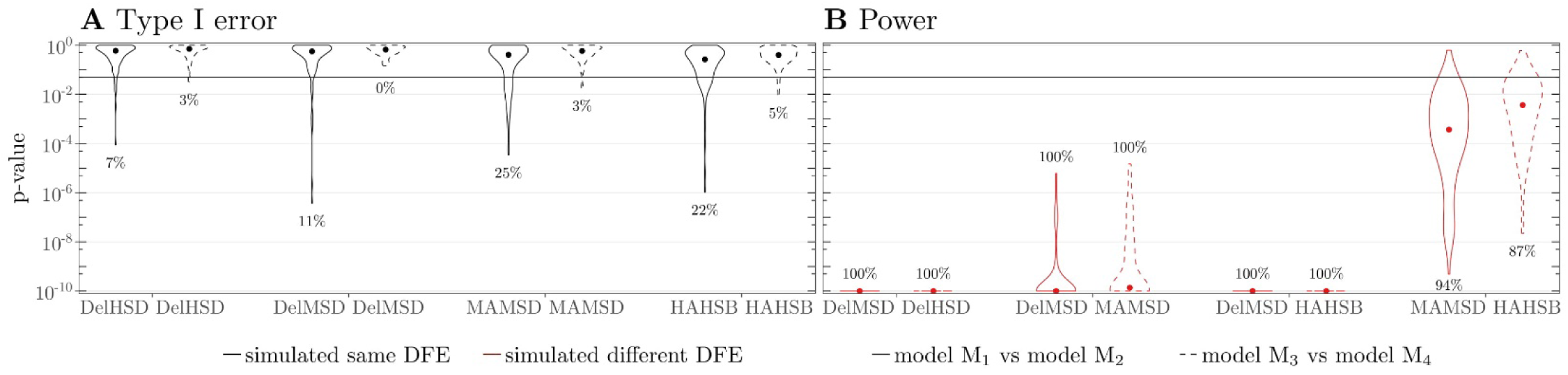
Violin plot of p-values for LRTs for different DFEs across SFS when data is simulated under various DFEs. DFE types used for simulations are indicated on the x-axis and detailed in Table S1. The LRT is performed using χ^2^ distribution and the maximum likelihoods found when inferring the parameters under models M_1_ and M_2_ (solid line) or models M_3_ and M_4_ (dashed line), described in Table 1 in the main text. The LRT is performed using datasets containing approximately 10MB of sites. (A) Type I error: polyDFEv2.0 was ran on 2 datasets simulated using the same DFE (black outline). (B) Power: polyDFEv2.0 was ran on 2 datasets simulated using different DFEs (red outline). Dots indicate the median. The horizontal lines show the 5% threshold and numbers in the plot indicate the percentage of p-values that are under the 5% threshold. For visualization reasons, p-values smaller than 10^−10^ have been changed to 10^−10^.

**Figure S2.**
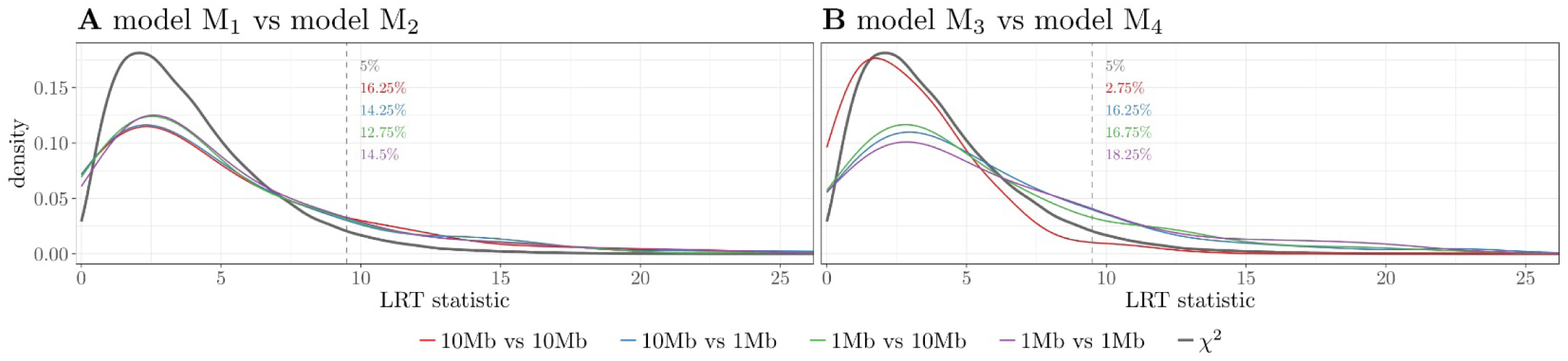
Empirical null distribution of the LRT statistic used for the LRT. The distribution is obtained from the maximum likelihoods found when inferring the parameters under models M_1_ and M_2_ (A) or models M_3_ and M_4_ (B) (described in Table 1 in the main text) from datasets that were simulated using the same DFE (Table S1). Using datasets of different size (1 or 10 Mb), we obtain different empirical distributions (shown in different colors). The χ^2^ distribution with 4 degrees of freedom, which is expected to be asymptotically a good approximation for the LRT, is given in gray. The vertical dotted line shows the 5% cutoff from the χ^2^ distribution, while the numbers in the plot indicate the percentage of LRT statistics that are above this asymptotic cutoff, for each empirical distribution.

**Figure S3.**
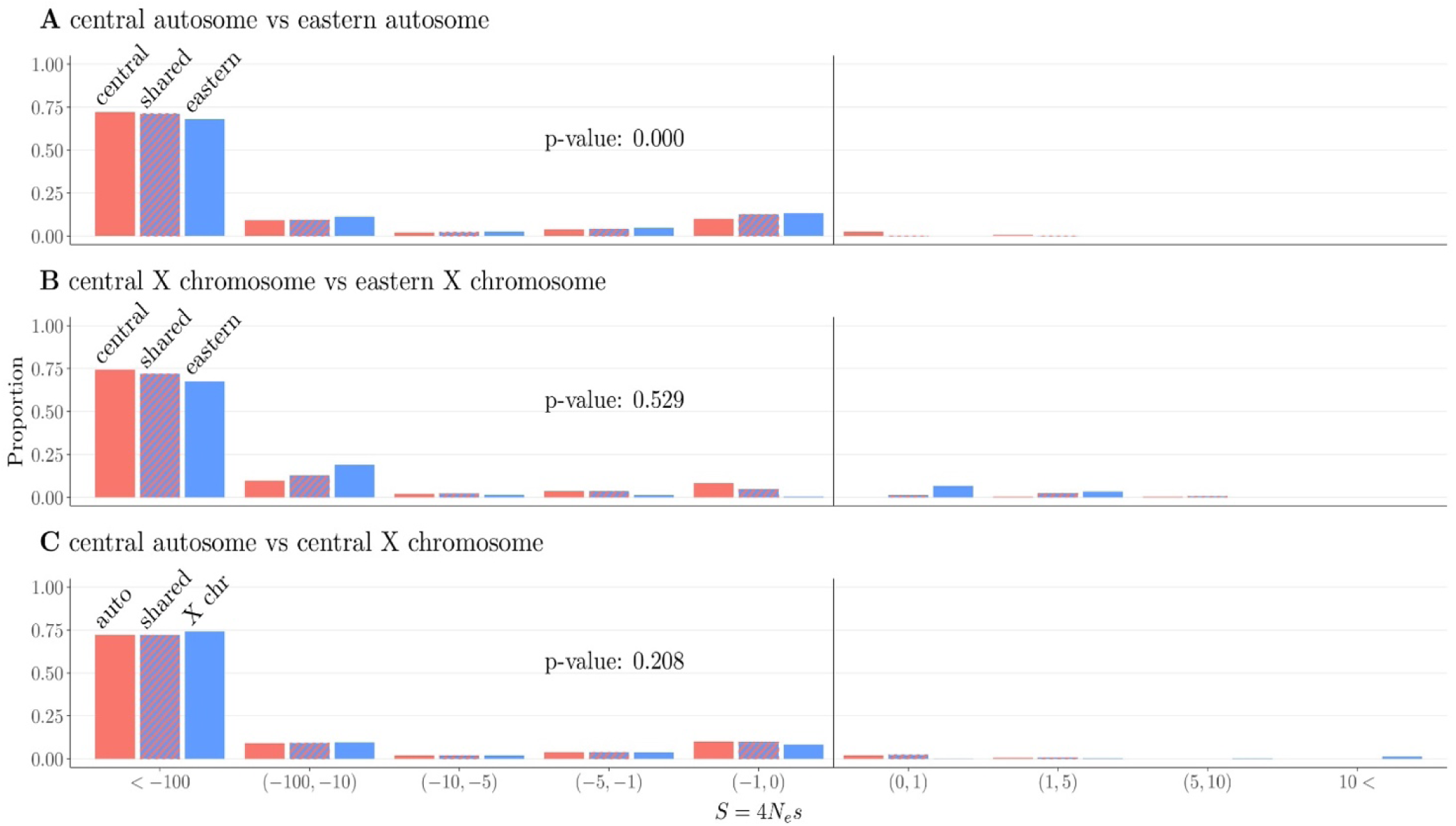
Inferred DFE on the chimpanzee SFS datasets. The DFE model fitted comprises both deleterious mutations (negative scaled selection coefficients ***S***) and beneficial mutation effects (positive **S**). The DFE was either inferred under model M_4_, fitting DFE parameters independently for each dataset (red and blue), or under model M_3_, assuming shared DFE parameters for both datasets (hashed red and blue). The p-values were obtained from LRTs comparing models M_3_ and M_4_ using the χ^2^ with 4 degrees of freedom as null distribution. Note: Inference was made with a continuous DFE (mixture of Gamma and exponential) but for graphical convenience DFE are discretized in classes of scaled selection coefficients

**Figure S4.**
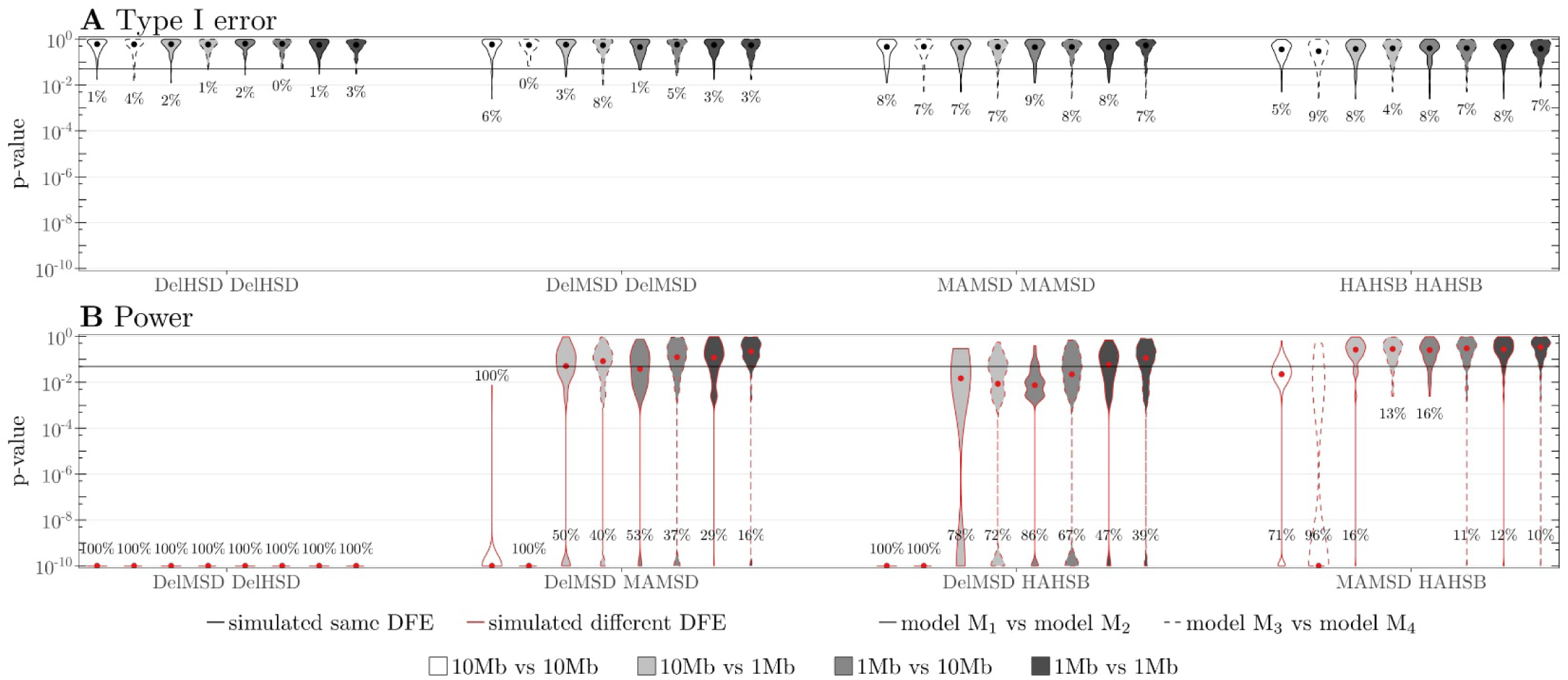
Violin plot of p-values for LRTs for evidence of different DFEs in the SFS on data simulated under various DFEs, given on the x-axis and detailed in Table S1. The LRT is performed using the empirical null distribution (Figure S2) and the maximum likelihoods found when inferring the parameters under models M_1_ and M_2_ (solid line) or models M_3_ and M_4_ (dashed line), described in Table 1 in the main text. The LRT is performed using datasets of different sizes. (A) Type I error: polyDFEv2.0 was ran on 2 datasets simulated using the same DFE (black outline). (B) Power: polyDFEv2.0 was ran on 2 datasets simulated using different DFEs (red outline). Dots indicate the median. The horizontal lines show the 5% threshold and numbers in the plot indicate the percentage of p-values that are under the 5% threshold. For visualization reasons, p-values smaller than 10^−10^ have been set to 10^−10^.

